# Prefrontal Cortical Deficits are a Putative Susceptibility Factor for PTSD

**DOI:** 10.1101/2024.10.18.619047

**Authors:** Rebecca Nalloor, Khadijah Shanazz, Almira Vazdarjanova

## Abstract

A subset of people who experience a traumatic event develop Post-Traumatic Stress Disorder (PTSD) suggesting that there are susceptibility factors influencing PTSD pathophysiology. While post trauma sequelae factors are extensively studied, susceptibility factors are difficult to study and therefore poorly understood. To address this gap, we previously developed an animal model - Revealing Individual Susceptibility to PTSD-like phenotype (RISP). RISP allows studying susceptibility factors by identifying, before trauma, rats that are likely to develop a PTSD-like phenotype after trauma. Hypofunctioning prefrontal cortex has been reported in people with PTSD, however, it is unclear if it is a susceptibility factor, sequalae factor, or both. Using RISP, we tested the hypothesis that altered medial prefrontal cortical (mPFC) function is a putative susceptibility factor. Susceptible male rats showed altered expression of plasticity-related immediate early genes (*Arc* and *Homer1a*) in the Prelimbic and Infralimbic subregions of the mPFC following spatial learning. Susceptible rats also showed deficits in attentional set shifting task when task demands increased. These findings suggest that Susceptible rats have mPFC deficits both at the cellular and functional level before trauma. Consistent with the findings in rats, military personnel who showed pre-trauma deficits in cognitive tasks involving mPFC developed PTSD post-trauma. Combined, these findings suggest that mPFC deficits are a putative susceptibility factor for PTSD and enhancing mPFC function in susceptible individuals before trauma may confer resilience to developing PTSD.

**Summary:** A subset of people who experience a traumatic event develop Post-Traumatic Stress Disorder (PTSD) suggesting that there are susceptibility factors influencing (PTSD) pathophysiology. While post trauma sequelae factors are extensively studied, susceptibility factors are difficult to study and therefore poorly understood. In the manuscript, we report data from both an animal model of susceptibility to a PTSD-like phenotype, and from military personnel, which is consistent with the hypothesis that deficits in prefrontal cortex function exist in susceptible individuals before PTSD-inducing trauma. The reported experiments include both a cellular molecular technique (*Arc/Homer 1a* catFISH), based on the expression of effector immediate-early genes necessary for consolidation of learning-induced plasticity, as well as behavioral experiments for assessing cognitive function in rats and humans.

## Introduction

Post-Traumatic Stress Disorder (PTSD) is a complex mental disorder that develops in a subset of people exposed to a traumatic event and results in long-lasting emotional and functional impairments (1). Individual susceptibility to developing PTSD has long been acknowledged (2– 5), yet it is poorly understood. To facilitate investigations into individual susceptibility, we have previously postulated a unifying framework of risk, susceptibility and sequelae factors. (6)

We define susceptibility factors as those aspects that are present and can be identified before trauma and manipulated such that they affect the onset and progression of PTSD after trauma (6). To identify and study susceptibility factors, we developed an animal model, RISP, (Revealing Individual Susceptibility to PTSD-like phenotype). Using RISP, one can classify male rats before trauma as Resilient or Susceptible to developing a PTSD-like phenotype based on a priori set of criteria of acoustic startle response and anxiety-like behaviors four days after exposure to a mild stressor. Susceptible, but not Resilient, rats subsequently displayed a PTSD-like phenotype with impaired fear extinction and sustained increase in acoustic startle response after a traumatic event (7).

Using the RISP model in conjunction with a sensitive cellular imaging technique called *Arc/Homer1a* catFISH (cellular compartment analysis of temporal activity by fluorescent in-situ hybridization using *Arc* and *Homer 1a*) we previously reported a putative susceptibility factor-impaired hippocampal learning-induced plasticity before trauma (8). We demonstrated that Susceptible, compared to Resilient, rats have impaired expression of immediate-early genes (IEGs) in the ventral/temporal and altered pattern of IEG expression in the dorsal/septal hippocampus during spontaneous spatial learning, a relatively low stress event. These differences were detected before the animals experienced a traumatic event suggesting that altered hippocampal function is a putative susceptibility factor. Such findings are consistent with reports in humans where hippocampal structural and functional deficits before trauma have been associated with increased severity of PTSD (5,9–11).

Other brain regions functionally related to the hippocampus such as the medial prefrontal cortex (mPFC) and amygdala have also been shown to have structural and functional alterations in people with PTSD (12–15). It is unclear however, if these alterations exist before trauma and thus contribute to the onset of PTSD or develop because of the trauma. We hypothesized that mPFC deficits are also a putative susceptibility factor for PTSD and can be detected before trauma. We tested this hypothesis by examining learning-induced plasticity in the Prelimbic (PL) and Infralimbic (IL) regions of the mPFC in Susceptible and Resilient rats before trauma using *Arc/Homer 1a* catFISH (16,17).

*Arc* (activity-regulated cytoskeleton-associated protein) and *Homer1a* (*H1a*) are effector immediate-early genes (IEGs) that are very low at baseline and are coordinately expressed during learning (16,18–21). They are reliable markers of plasticity, not solely neuronal activity (22,23) and are necessary for memory consolidation (20,24–26). Furthermore, *Arc* expression reliably predicts the pattern of plasticity assessed electrophysiologically a day after learning (22,23). Additionally, *Arc* and *H1a* mRNA have different lengths such that, using a full-length Arc probe and a similar sized probe targeting the 3’ UTR of *H1a*, it is possible to dissociate neuronal ensembles that have initiated plasticity during two events experienced 20-30 mins apart. Neurons with *H1a* foci of transcription mark those responding to the first event and the ones with *Arc* foci mark neurons responding to the second event. Neurons with both *H1a* and *Arc* foci have responded to both events (8,16,27,28)(Fig 1B). One advantage of *Arc/H1a* catFISH is that the dissociation of the neuronal ensembles responding to these events can be done across multiple brain regions in the same animals. Combined with the RISP model, *Arc/H1a* catFISH allowed us to examine whether relative plasticity across different brain regions is altered in Susceptible compared to Resilient rats.

**Figure 1.**
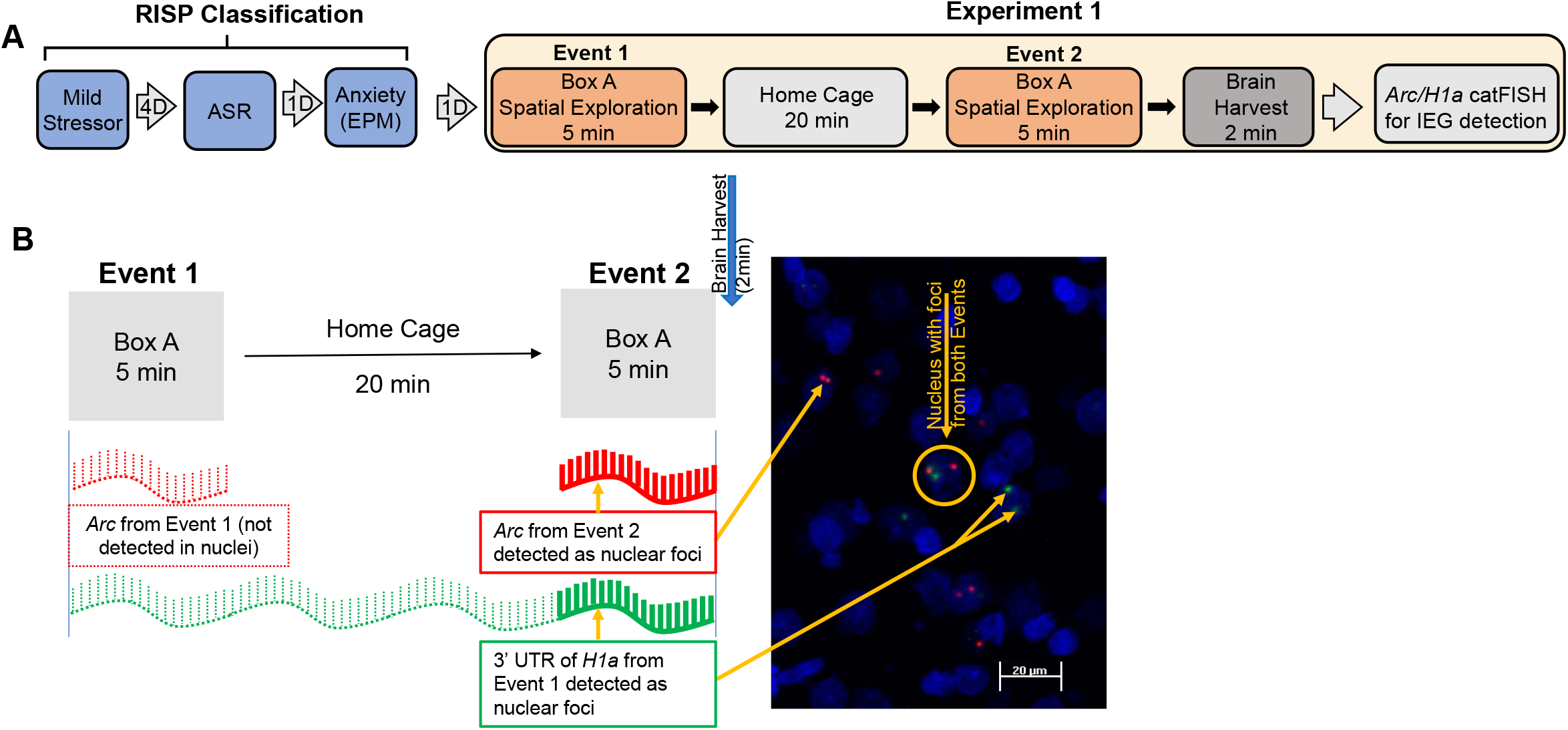
A) RISP classification and design of Experiment 1. B) Schematic representation of *Arc/Homer1a* catFISH with a representative image from IL– Nuclei (blue), *Arc* (red), *H1a* (green)

**Figure 2.**
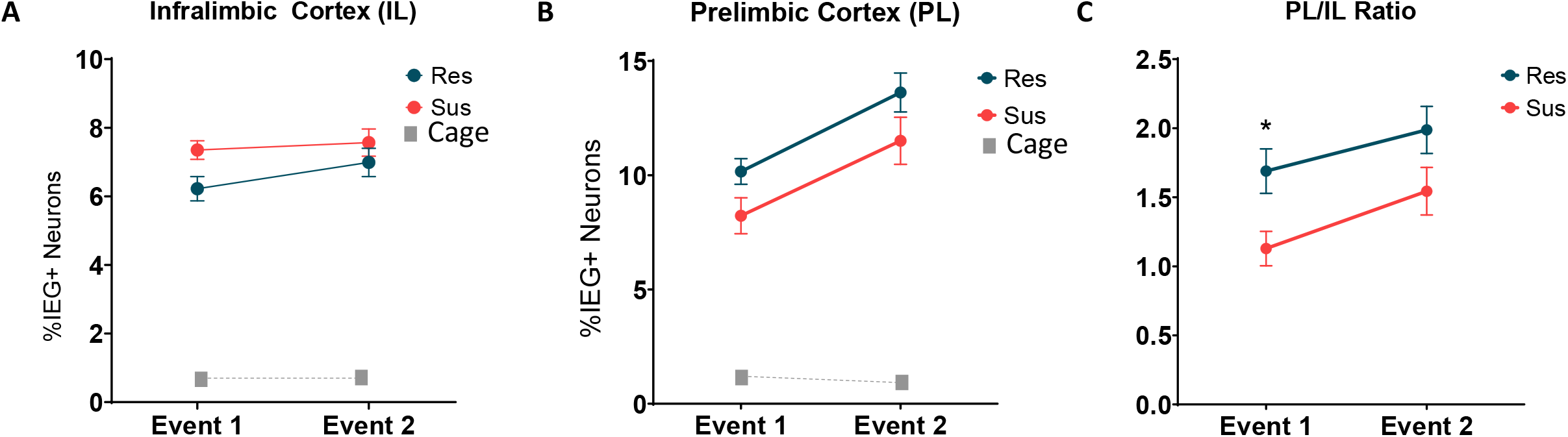
*Arc* and *H1a* expression in the mPFC of Resilient (Res, n=8) and Susceptible (Sus, n=6) rats during Event 1 and Event 2 in the A) IL, B) PL, and C) The Pl/IL ratio of activated ensembles. *p = 0.024 (Fisher’s LSD after RM-ANOVA)

We also hypothesized that any differences in plasticity between Resilient and Susceptible rats will also be detected functionally with appropriate cognitive tests. Attentional Set-Shifting is a task that is commonly used to test cognitive flexibility in animals (29,30). Impairments in this task are associated with structural changes and functional deficits in the mPFC (31–33). Therefore, we tested whether the Resilient and Susceptible rats differ on their performance on the Attentional Set-Shifting task.

To begin evaluating the translational relevance of these hypotheses, we compared pre-deployment performance on a battery of cognitive tests of military personnel who developed post-deployment PTSD and those who did not. We hypothesized that susceptibility factors also exist in humans and can be functionally detected before a PTSD-inducing trauma.

## Materials and Methods

### Subjects

#### Animal

Two-three-month-old, adult (250-300g) male Sprague-Dawley rats (Charles River Laboratories Inc, Wilmington, MA) were double housed on a 12 hrs light/dark cycle (lights on at 7:00 am) with food and water freely available except during Experiment 2 (Attentional set shifting) when they were food restricted (Fig 3A). Testing was performed between 9:00 am and 5:00 pm.

**Figure 3.**
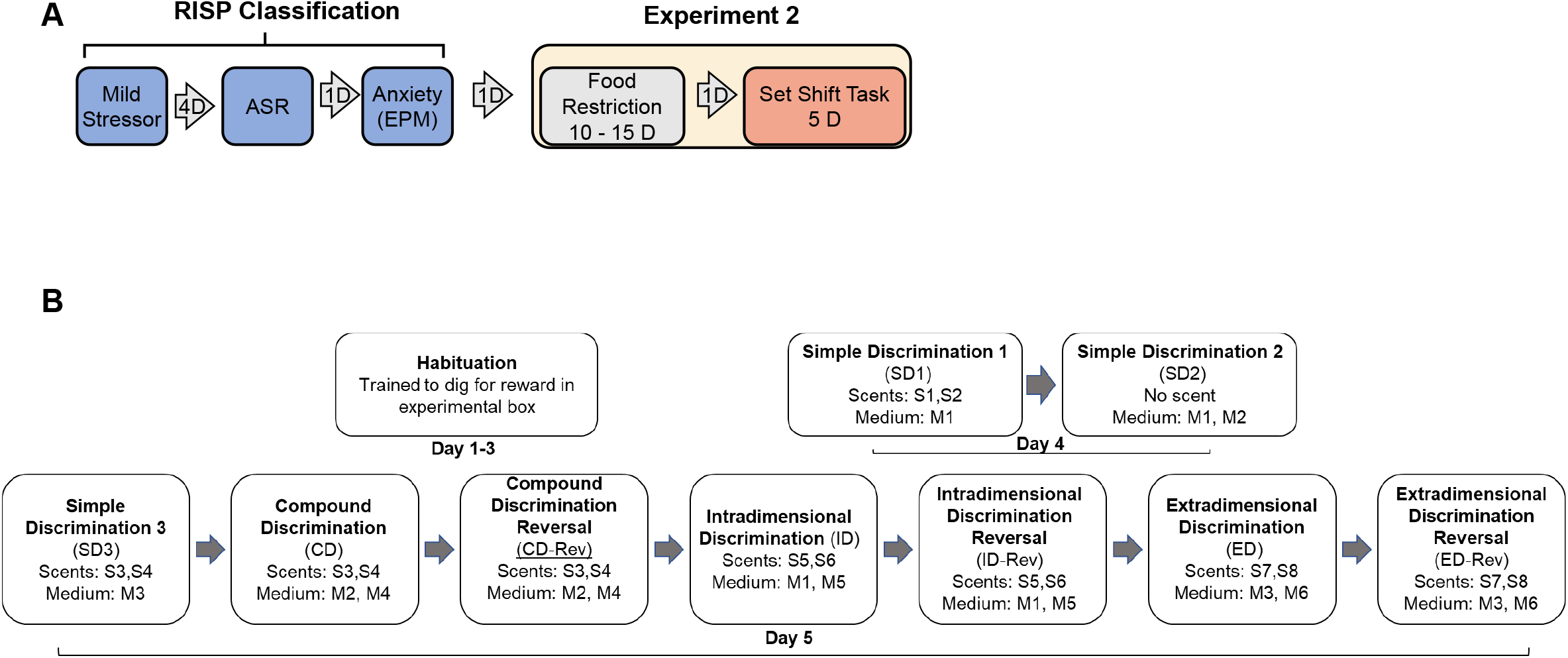
A) RISP classification and design of Experiment 2 B) Details of the Attentional Set-Shifting task

#### Human

De-identified pre-deployment ANAM (Vista Life Sciences, Inc. Englewood, CO) data and post deployment PTSD status from a small group of male (age: 23-58 yrs) active-duty military personnel was obtained (Fort Gordon, Augusta, GA).

### Rodent behavioral procedures

The number of rats per group is listed under each experiment and in each figure. All rodents were handled for 2–3 minutes for three consecutive days, starting three days after arrival. Behavioral testing began after handling. All behavioral procedures were approved by the Institutional Animal Care and Use Committee (IACUC), Augusta University and Charlie Norwood VA Medical Center, Augusta, GA.

### RISP Animal Model

The RISP (Revealing Individual Susceptibility to a PTSD-like Phenotype) animal model of PTSD is described in detail in Nalloor, 2011 and Alexander 2020 (6,7). Briefly, four days after a brief exposure to a mild stressor (cat hair), rats were tested for their acoustic startle response (ASR) and anxiety-like behavior on the elevated plus maze (EPM). Using a priori set criteria, they were classified as:

1. Resilient (Res) - low startle (average ASR and more than 7 of 15 individual ASR smaller than the group average ASR) and low anxiety-like behavior (at least 1 entry into the open arm of the EPM)
2. Susceptible (Sus) - high startle (average ASR and 6 or more of 15 individual ASR greater than the group average ASR) and high anxiety-like behavior (no entries into the open arms of the EPM).
3. Intermediate-Animals meeting neither set of criteria; excluded from further analysis.

## Experiment 1

The design of Experiment 1 is summarized in Figure 1A. The behavior of these animals during classification was previously reported in Nalloor et al 2014 (8). Rats classified as Resilient (n = 8) or Susceptible (n= 6) were allowed to explore a novel box (50cm x 10cm x 19cm) twice for 5 mins, 20 mins apart (Event 1 and Event 2, respectively). The rats were in their home cages during the 20 min between the Events. Fear behavior and locomotion were assessed. Freezing, a measure of fear behavior, was defined as lack of all movement except for respiration. Number of Crossings, a measure of locomotor activity, was defined as all four paws and the base of the tail crossing over the midline of the box. Immediately after Event 2, rats were quickly anesthetized (<1.5 min) and decapitated. Caged control rats were removed from the cages either before the first or second behavioral event and were immediately anesthetized for brain harvesting and assessing of baseline *Arc/H1a* expression.

### Brain harvesting

The brains were rapidly harvested and flash-frozen in 2-methyl butane (time from initiation of anesthesia to freezing of the brain < 3 min). They were stored at -80°C until further processing.

### *Arc/Homer1a* catFISH

The right hemisphere of brains from Res and Sus rats were blocked together in freezing medium with a maximum of 6 brains per block. Twenty-micron thick coronal sections were cut on a cryostat and mounted on glass slides. The slides were stored at -20°C or -80°C until further processing. The slides were processed for *Arc/Homer1a* catFISH as described by (8,27,34,35) to detect transcription foci of *Arc* and *Homer1a (H1a)*. Briefly, after fixing in 4% paraformaldehyde and permeabilizing the tissue in 1:1 acetone/methanol, a fluorescein-labeled full-length *Arc* and digoxigenin-labeled *H1a* antisense riboprobe, targeting the 3’ UTR of *H1a* mRNA, were applied and the slides incubated overnight at 56°C. Following hybridization, slides underwent stringency washes, endogenous peroxidase quenching and blocking for non-specific antibody binding. The digoxigenin tag was detected sequentially with horseradish peroxidase-conjugated anti-digoxigenin antibody (Fab fragments, Roche Diagnostics, Indianapolis, IN) and tyramide amplification reaction with SuperGlo Green (Fluorescent Solutions, Augusta, GA). Following a second peroxidase quenching step, the fluorescein tag was revealed with horseradish peroxidase-conjugated anti-fluorescein antibody (Roche) and tyramide amplification reaction with SuperGlo Red (Fluorescent Solutions).

Riboprobes were generated using commercial transcription kits (Ambion MaxiScript, Life Technologies, Grand Island, NY; or AmpliScribe; Epicentre Biotechnologies, Madison, WI) and digoxigenin-UTP or fluorescein-UTP RNA labeling mixes (Roche). Coverslips were mounted and nuclei were counterstained using Vectashield Mounting Medium with DAPI (Vector Laboratories, Burlingame, CA).

### Image Acquisition and Stereological Analysis

Mosaic (850×650µm) image stacks from Prelimbic (PL) and infralimbic (IL) regions of the medial prefrontal cortex (mPFC) were collected from at least 2 different slides per animal using a 20x objective on a Zeiss AxioImager/Apotome system. The mPFC (3.72-2.76 mm anterior to Bregma) was identified based on Figures 9-12 of The Rat Brain by Paxinos and Watson (36).

Unbiased stereological cell counting and classification were performed as follows: 1) Neuron-like cells in the regions of interest were segmented by using an optical dissector method (37) with the gene channels turned off to prevent selection bias; 2) segmented neurons were classified using Zeiss AxioVision imaging software. Putative glial cells (those with small, intensely, and uniformly stained nuclei) were excluded from the analysis. With the gene channels turned on, segmented neurons were classified as *Arc*+, *H1a*+ or *Arc/H1a*+ depending on whether they had foci of transcription for *Arc, H1a*, or both, respectively. Foci were quantified only when they were visible on 3 or more consecutive planes. Cells without any foci were classified as Negative. For purposes of clarity, we refer to all *H1a+* cells as an ‘activated ensemble’ that initiated plasticity-related IEG expression during Event 1, and all *Arc+* cells as an activated ensemble from Event 2.

A schematic representation of the time course of Events in Experiment 1 and the corresponding expression of *Arc* and *H1a* are shown in Figure 1B.

## Experiment 2

The experimental design is summarized in Figure 3A. Similar to Experiment 1, rats were classified using the RISP model. Resilient (n =11) and Susceptible (n = 9) rats were food restricted to maintain their body weight at ∼ 85% of pre-restriction weight.

Attentional Set-Shifting task: Details of the order of discrimination tasks are provided in Figure 3B.

### Habituation

On Days 1-3, rats were habituated to a 30 × 25 × 30 cm chamber containing 2 cups (5 cm diam. X 4 cm deep) inserted into the floor. They were trained to dig for a buried food reward (1/2 of a Froot Loops cereal piece) in various scented and unscented media placed into the cups. The animals were exposed to all scents and media.

### Training and testing

On Day 4 and 5, each animal was placed into the chamber for up to 1 min per trial and allowed to dig in one of the two cups for the food reward (1/4 of a Froot Loops piece). One of the two cups (left or right) was baited with a food reward, the other cup was not; trials were scored as “correct” if the animal chose to dig in the baited cup. After retrieving the food reward or after they stopped digging in the incorrect cup, animals were removed and placed into a holding cage between trials. Each discrimination task was complete when the animal performed 6 consecutive correct trials. The length of each trial and number of trials to reach criterion were recorded. The presentation of the reward was pseudorandomized for right/left placement and medium/scent combination. The list of media and scents is provided in Table 1.{Insert Table 1}

**Table 1:** Set Shifting Media and Scents.

|  |  |  |  |  |  |  |  |  |
| --- | --- | --- | --- | --- | --- | --- | --- | --- |
| Media | M1 | M2 | M3 | M4 | M5 | M6 |  |  |
|  | Corn cob bedding | Paper pellet bedding | Wood shavings | Sawdust | Plastic beads | Sand |  |  |
| Scents | S1 | S2 | S3 | S4 | S5 | S6 | S7 | S8 |
|  | Basil | Nutmeg | Coffee | Cinnamon | Mint | Cocoa | Cumin | Oregano |

## Human Data

The timepoints of collection of the deidentified human data are provided in Figure 5A. The pre-trauma cognitive assessment was performed using ANAM (Automated Neuropsychological Assessment Metrics, Vista Life Sciences, CO), a battery of computerized cognitive tests that also evaluates simple reaction time at the first and second half of the battery. The ANAM data was collected before combat exposure. At this time, none of the subjects met the criteria for PTSD diagnosis. Subsequently, all subjects encountered severe combat related traumatic events that resulted in TBI. Following this trauma, some developed PTSD (PTSD; n=3) and some did not (NoPTSD; n=6). The PTSD evaluation and diagnosis were performed by qualified clinicians at Eisenhower Army Medical Center, Augusta, GA.

Both NoPTSD and PTSD groups were compared on the following outputs of their pre-deployment ANAM: Simple Reaction Times, Accuracy, Throughput and Reaction Time for each cognitive test. In order to normalize for possible differences in individual reaction time, we calculated Cognitive Processing Time:

Cognitive Processing Time = Reaction Time for test – (Simple Reaction Time 1+ Simple Reaction Time 2)/2

### Statistical Analyses

Two-way repeated measures ANOVA with Group and Event (Fig. 2) or Task (Fig. 4) as independent variables was used to compare IEG expressing ensembles or number of trails to criterion, respectively. Statistically significant main effects were followed up by a one-way ANOVA for within group comparison. The differences in ensemble size within each group between Event 2 and Event 1 (Event 2/Event1) were evaluated using one group t-test with a hypothesized difference of 1. Student t-tests were used to compare performance of No PTSD and PTSD groups in the different cognitive tests (Fig. 5B-D) (GraphPad 10). Differences were considered statistically significant at p< 0.05.

**Figure 4.**
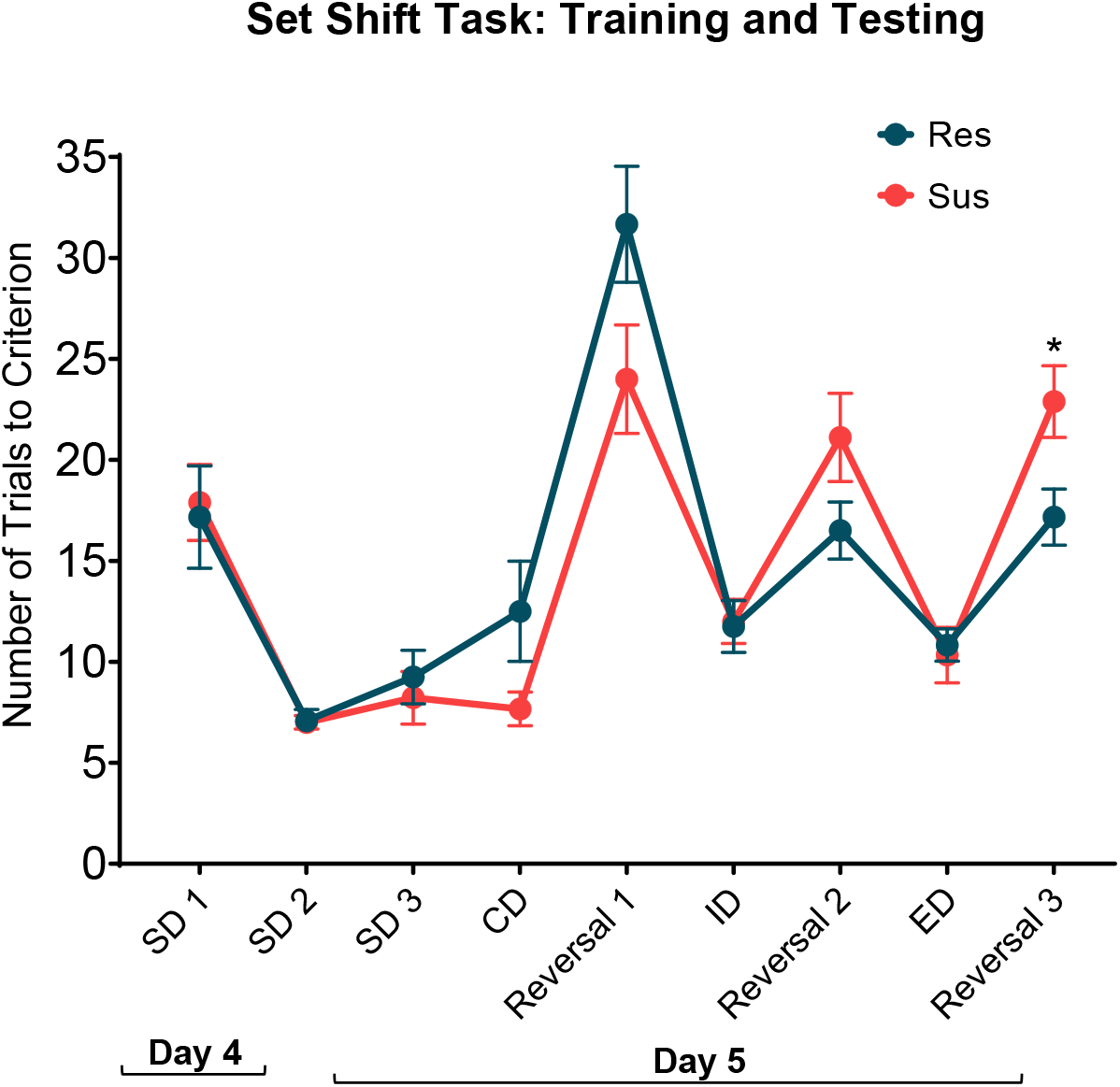
Performance of Resilient (Res, n=11) and Susceptible (Sus, n=9) rats on the training (Day 4) and testing day (Day 5) of Attentional Set-Shifting task. RM-ANOVA of the number of trials to criterion for each of the discriminations. SD-Simple Discrimination, CD-Compound Discrimination, ID-Intradimensional Discrimination, ED-Extradimensional Discrimination. *p = 0.022 (Fisher’s LSD post hoc)

**Figure 5.**
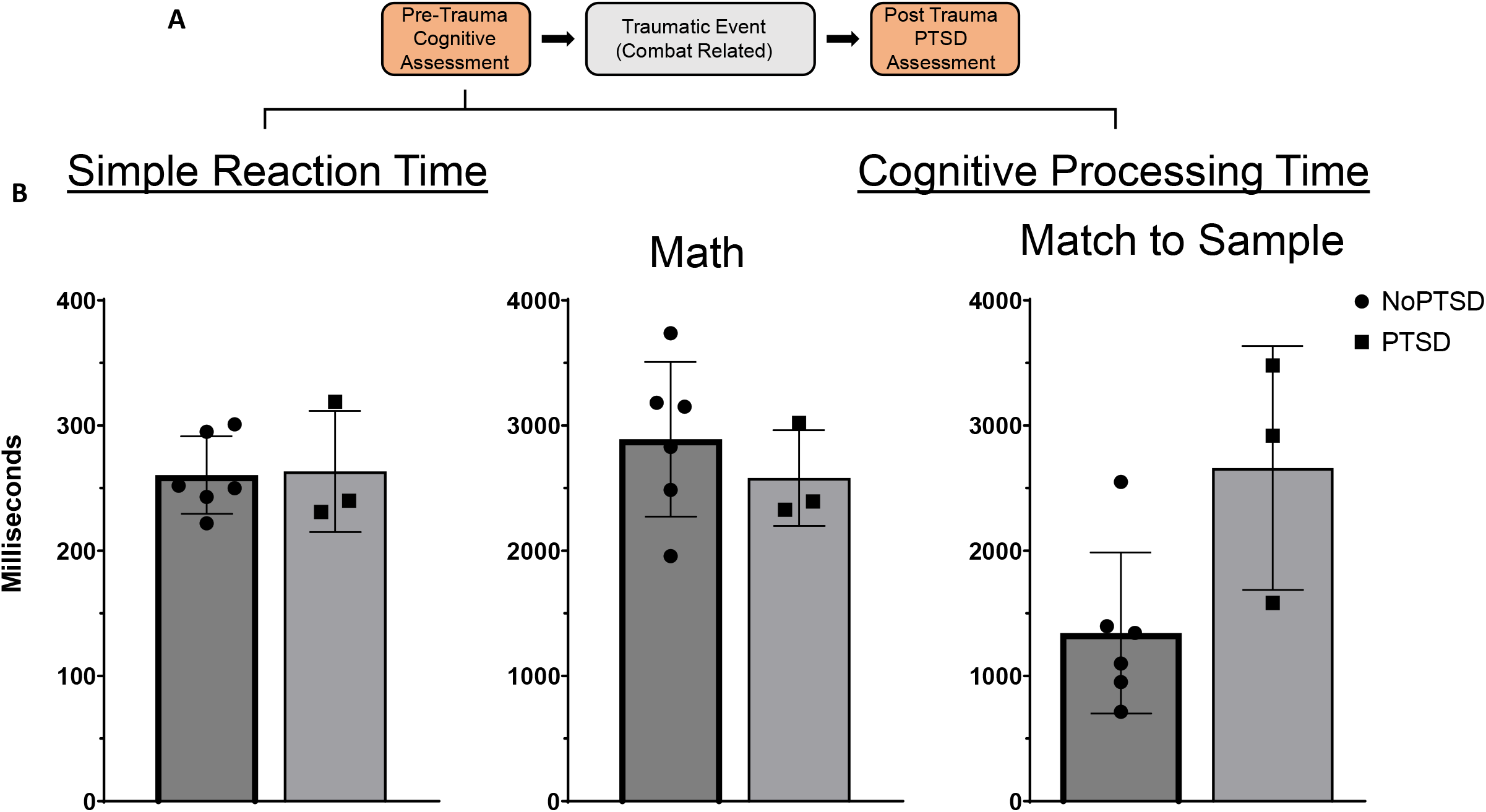
Pre-deployment cognitive performance of military personnel with post-deployment PTSD status A) Timeline of key events reflecting the point at which the presented data was collected; B - D) Pre-trauma ANAM data. *p= 0.042.

## Results

### Resilient and Susceptible rats have different patterns of *Arc/Homer1a* expression in the mPFC in response to spatial exploration

Fig. 1A illustrates the RISP model and design of Experiment 1. The behavior of these animals during classification was previously reported in Nalloor et al 2014. Here, one of the Resilient rats from the original data set was excluded due to damage to the mPFC during tissue processing. The *Arc/Homer1a* results reported here are from eight Resilient and six Susceptible rats.

During the two spatial exploration events (Event 1 and Event 2), all animals explored the box well. There were no differences between Resilient and Susceptible rats in the Number of Crossings (Group effect: F(1,12)= 0.052, p= 0.823, β= 0.055 and no Group x Event interaction: F(1,12)= 2.260, p= 0.159, β= 0.271). None of the rats displayed freezing > 5%. Despite similar behavior, differences were detected between groups in mPFC IEG ensembles responding to these events (Fig. 2A-C). This data is based on an average cell count of 675 per animal/per brain region.

In IL (Fig 2A), there was an overall increase in ensemble size during Event 2 (Event effect: F (1,12) = 6.212, p = 0.028, β = 0.628 with no Group effect: F (1,12) = 2.356, p=0.151, β= 0.281 and no Group x Event interaction: F (1,12) = 1.955, p=0.187, β= 0.240). This ensemble size difference was driven by the Res group as the ratio of Event 2/ Event 1 was significantly different from 1 only for the Res group (p=0.026; Sus: p = 0.518). Baseline *Arc* and *H1a* expression in caged controls was <1%.

Similarly, in PL (Fig 2B), there was an increase in ensemble size during Event 2 (Event effect: F (1,12) = 28.158, p = 0.0002, β = 0.999). There was also a nearly significant Group effect (F (1,12) = 4.478, p = 0.056, β = 0.485) with no Group x Event interaction (F (1,12) = 0.019, p = 0.892, β = 0.052). Furthermore, both groups showed increased ensemble size: Event 2/Event 1 was significantly > 1 for both Res and Sus groups (p < 0.0001). Baseline *Arc* and *H1a* expression in caged controls was <1%.

We further evaluated this difference in the pattern of IEG expression between PL and IL in Res and Sus rats by comparing their PL/IL ratio. The PL/IL ratio increased for both groups during the second exploration (Event effect: F (1,12) = 10.13, p= 0.008, β = 0.844). This increase was different between groups such that the PL/IL ratio was significantly higher in the Res rats compared to the Sus rats (Group effect: F (1,12) = 6.11, p= 0.029, β = 0.621) with no Group x Event interaction (F (1,12) = 0.27, p=0.610, β = 0.076). Res rats had a bigger PL ensemble size than IL during both Event 1 (PL/IL = 1.69, p = 0.004) and Event 2 (PL/IL = 1.99, p = 0.0007), while in Susceptible rats this was true only for Event 2 (PL/IL= 1.54, p = 0.025). In Sus rats PL and IL ensemble sizes were similar during Event 1 (PL/IL = 1.13, p = 0.349) (Fig 2 C). Furthermore, during Event 1 PL/IL ratio was significantly higher in Res compared to Sus rats (p = 0.024). Combined, these data suggest that in Resilient rats, the PL consistently engages a larger ensemble than IL while this PL dominance is not seen in the Susceptible rats.

It is important to stress that these differences were detected in animals who had not experienced trauma. Overall, these findings demonstrate that there were differences in the pattern of learning-induced plasticity in the mPFC between Resilient and Susceptible rats before trauma.

### Susceptible rats have behavioral deficits under high cognitive demand

We examined if the above described differences in the pattern of plasticity have functional correlates by comparing behavior of Resilient and Susceptible rats in a mPFC dependent task. Figure 3A shows the RISP model and the design of Experiment 2.

As seen in Figure 4, The overall pattern of performance during the attentional set-shifting task was different between groups (Group x Discrimination interaction: F (8, 144) = 2.94, p< 0.005, β = 0.951) without an overall Group effect F (1, 18)= 0.67, p= 0.423, β = 0.117). Specifically, Resilient rats showed improvement in the reversal tasks as the training progressed, while Susceptible rats did not. Notably, Susceptible rats had significantly higher number of trails to criterion compared to Resilient rats on Reversal 3 (p= 0.022). There were no differences in the performance during simple and complex discriminations as evidenced by similar number of trials to criterion between groups (SD1, p= 0.821; SD2, p= 0.899; SD3, p= 0.587; CD, p= 0.087; ID, p=0.884; ED, p=0.758). This shows that the Susceptible rats were able to learn and perform the simple versions of the task, but the deficits became evident under higher cognitive demand.

### Military personnel with combat-related PTSD showed altered performance in PFC dependent task at pre-deployment evaluation

As seen in figure 5D, soldiers who developed combat-related PTSD post-deployment had a higher Cognitive Processing Time on spatial Match to Sample task, a hippocampal and prefrontal cortex dependent task, during their pre-deployment evaluation (p=0.042). This difference was specific to the Match to Sample task as both groups had similar Cognitive Processing Time in other cognitive tasks such as Math (p=0.460) (Fig 5C). Both groups also had similar Simple Reaction Time (0.961) (Fig 5B). Furthermore, both groups had similar accuracy on all tasks (p> 0.98). The higher cognitive processing time in the Match to Sample task is consistent with functional deficits in the PFC and hippocampus and is independent of any individual differences in simple reaction time.

It is important to stress that all soldiers experienced severe combat-related trauma that resulted in TBI and significantly affected their post-trauma performance (not reported here as it is not relevant to the tested hypothesis).

## Discussion

The set of studies reported here aimed to test the hypothesis that mPFC deficits are a putative susceptibility factor for PTSD that can be detected before trauma in both rats and humans. The results provided evidence in support of this hypothesis at both the cellular and behavioral levels in rats and at the behavioral level in humans. At the cellular ensemble level, rats pre-classified according to the RISP model as Susceptible to developing a PTSD-like phenotype had a different pattern of plasticity-induced IEG expression in the mPFC than Resilient rats. On the behavioral level, Susceptible rats showed deficits under higher cognitive demand and people who later developed PTSD showed longer cognitive processing time in a PFC- and hippocampus-dependent task before trauma.

At the cellular level, the differences between Resilient and Susceptible rats were in the relative contribution of PL and IL in spontaneous spatial learning. All rats engaged PL and IL regions of the mPFC during spatial exploration as evidenced by the presence of above baseline number of neurons expressing plasticity-associated immediate-early genes (*Arc* and *H1a*) in both groups. However, there was a difference in the pattern of *Arc/H1a* expression between Resilient and Susceptible rats such that Resilient rats consistently engaged a larger ensemble in PL than IL while this PL dominance was not seen in the Susceptible rats. To our knowledge, this is the first study to show mPFC involvement in spontaneous spatial learning. The differences in plasticity between Resilient and Susceptible rats were present even when there were no differences in the behavior during the task. The fact that these differences are present before trauma suggests that these brain regions may respond differently to a traumatic event.

The PL and IL have distinct roles in cognition and emotional regulation (38,39). But reciprocal connections between these brain regions allow information exchange that leads to flexible and adaptive behavior (40,41). Both PL and IL also receive unidirectional projections from the hippocampus (42,43) and are implicated in fear learning and extinction (38,44,45). Our previous data show that in the same Susceptible animals the hippocampus (dorsal and ventral) has altered function compared to Resilient animals before trauma (8). Combined, this data suggests that there are multiple brain regions that are different between Resilient and Susceptible animals in information processing before trauma. Importantly, these differences are in brain regions that are well documented to have impaired functioning in people with established PTSD (14,46,47). Such findings expand our understanding of the pathophysiology of PTSD by separating susceptibility from sequalae factors.

It is important to note that the differences on the cellular level that are reported here (Experiment 1) were detectable because of the unique strengths offered by the *Arc/H1a* catFISH. Due to the restricted time of learning-induced *Arc/H1a* expression (<10 min) as detected by intranuclear foci of transcription, the temporal separation of the two events (30 min), and the very low level of baseline *Arc* and *H1a* expression (<1%), the activated neuronal ensembles are highly specific to each event. This specificity allows a more precise evaluation of changes in patterns of activation across multiple brain regions. The prominent differences in the PL and IL were not in the size of the ensembles induced by the two events but in the relative activation across the two brain regions. In addition, these findings are based on a large number of neurons per brain region and thus are representative of each region.

The differences at the cellular ensemble level were reflected at the behavioral level in a hippocampal/mPFC dependent task (Attentional Set Shifting) (32,48,49). While there were no differences in the initial learning of discrimination and reversal tasks, Susceptible rats showed impaired performance when cognitive load increased. The impaired performance developed gradually during the training with Susceptible rats starting off slightly better than Resilient in the Complex Discrimination and initial Intradimensional Reversal tasks but becoming slightly impaired in the second Intradimensional Reversal and then significantly impaired at the subsequent Extradimensional Reversal. Combined, these findings strongly suggest that the differences are underlined by altered mPFC function which guides flexible and adaptive behavior.

The behavioral findings demonstrated in rats paralleled those we observed in the human data set. Military personnel who developed PTSD post-trauma had cognitive deficits that were detectable pre-trauma. Similar to rats, these differences were not universal to all tasks but were specific to a hippocampus/mPFC-dependent task (spatial Match-To-Sample) (50–53). The differences in cognitive processing time were not based on motivation or general inability to understand or perform the tasks, as accuracy was not affected. They were also not due to individual differences in simple reaction time, as the latter was subtracted from the individual task reaction time for each participant to derive the Cognitive Processing Time. These considerations demonstrate the specificity of the reported findings. These findings demonstrate that pre-trauma functional differences exist and can be identified in humans as well.

In addition to the discussed strengths, the study has several limitations. First, although it would have been ideal to examine alterations in patterns of plasticity during the Attentional Set-Shifting task, this was not possible due to the temporal specificity of the *Arc/H1a* catFISH method: performing the task took much longer than the time window that is optimal for inducing and detecting *Arc/H1a* intranuclear foci of transcription (3-7 min). Second, the presented data is based on studies that included only male subjects. PTSD is a sexually dimorphic disorder, so the conclusions drawn may only be applicable to males. Additional studies in female subjects are required before generalizing findings to both sexes. Third, the human data presented here is preliminary as it is based on a small sample size. Furthermore, because all soldiers experienced TBI, it could have influenced the PTSD diagnosis. However, the human data findings are consistent with the animal data and thus warrant larger scale study in humans to examine cognitive determinants of susceptibility to PTSD with or without physical trauma.

Despite the stated limitations, combined, the findings suggest that mPFC deficits are a putative susceptibility factor for PTSD. Future studies will address whether enhancing mPFC function in susceptible individuals will confer resilience to developing PTSD. Building resilience is crucial for minimizing the number of people suffering from PTSD, given that it is difficult to treat, and treatments are resource intensive and benefit only a subpopulation of people suffering from PTSD.

## Acknowledgments

This material is based upon work supported in part by the Department of Veterans Affairs, Veterans Health Administration, Office of Research and Development, Biomedical Laboratory Research and Development, I01BX001978 and I01BX003890.

The authors thank Dr. Kristopher Bunting for technical support and Dr. Vaugh McCall for comments on a previous version of the manuscript. The contents of this manuscript do not represent the views of the VA or the United States Government.

## References

1. American Physiological Association. Diagnostic and statistical manual of mental disorders, 4th edition. Washington, DC: American Psychiatric Association; 2000.

2. Yehuda R, Bierer LM, Schmeidler J, Aferiat DH, Breslau I, Dolan S. Low Cortisol and Risk for PTSD in Adult Offspring of Holocaust Survivors. Am J Psychiatry. 2000 Aug 1;157(8):1252–9.

3. Childress JE, McDowell EJ, Dalai VVK, Bogale SR, Ramamurthy C, Jawaid A, et al. Hippocampal volumes in patients with chronic combat-related posttraumatic stress disorder: a systematic review. J Neuropsychiatry Clin Neurosci. 2013;25(1):12–25.

4. Logue MW, van Rooij SJH, Dennis EL, Davis SL, Hayes JP, Stevens JS, et al. Smaller Hippocampal Volume in Posttraumatic Stress Disorder: A Multisite ENIGMA-PGC Study: Subcortical Volumetry Results From Posttraumatic Stress Disorder Consortia. Biol Psychiatry. 2018 01;83(3):244–53.

5. Gilbertson MW, Shenton ME, Ciszewski A, Kasai K, Lasko NB, Orr SP, et al. Smaller hippocampal volume predicts pathologic vulnerability to psychological trauma. Nat Neurosci. 2002 Nov;5(11):1242–7.

6. Alexander KS, Nalloor R, Bunting KM, Vazdarjanova A. Investigating Individual Pre-trauma Susceptibility to a PTSD-Like Phenotype in Animals. Front Syst Neurosci. 2020;13.

7. Nalloor R, Bunting K, Vazdarjanova A. Predicting Impaired Extinction of Traumatic Memory and Elevated Startle. PLOS ONE. 2011;6(5):e19760.

8. Nalloor R, Bunting KM, Vazdarjanova A. Altered Hippocampal Function Before Emotional Trauma in Rats Susceptible to PTSD-Like Behaviors. Neurobiol Learn Mem. 2014 Jul;112:158– 67.

9. Gilbertson MW, Paulus LA, Williston SK, Gurvits TV, Lasko NB, Pitman RK, et al. Neurocognitive function in monozygotic twins discordant for combat exposure: relationship to posttraumatic stress disorder. J Abnorm Psychol. 2006;115:484–95.

10. Hayes JP, LaBar KS, McCarthy G, Selgrade E, Nasser J, Dolcos F, et al. Reduced hippocampal and amygdala activity predicts memory distortions for trauma reminders in combat-related PTSD. J Psychiatr Res. 2011;45:660–9.

11. Xie H, Claycomb Erwin M, Elhai JD, Wall JT, Tamburrino MB, Brickman KR, et al. Relationship of Hippocampal Volumes and Posttraumatic Stress Disorder Symptoms Over Early Posttrauma Periods. Biol Psychiatry Cogn Neurosci Neuroimaging. 2018 Nov 1;3(11):968–75.

12. Milad MR, Pitman RK, Ellis CB, Gold AL, Shin LM, Lasko NB, et al. Neurobiological basis of failure to recall extinction memory in posttraumatic stress disorder. Biol Psychiatry. 2009;66:1075–82.

13. Pitman RK, Rasmusson AM, Koenen KC, Shin LM, Orr SP, Gilbertson MW, et al. Biological studies of post-traumatic stress disorder. Nat Rev Neurosci. 2012 Nov 1;13(11):769–87.

14. Harnett NG, Goodman AM, Knight DC. PTSD-related neuroimaging abnormalities in brain function, structure, and biochemistry. Exp Neurol. 2020 Aug 1;330:113331.

15. Xiao S, Yang Z, Su T, Gong J, Huang L, Wang Y. Functional and structural brain abnormalities in posttraumatic stress disorder: A multimodal meta-analysis of neuroimaging studies. J Psychiatr Res. 2022 Nov 1;155:153–62.

16. Vazdarjanova A, McNaughton BL, Barnes CA, Worley PF, Guzowski JF. Experience-Dependent Coincident Expression of the Effector Immediate-Early Genes Arc and Homer 1a in Hippocampal and Neocortical Neuronal Networks. J Neurosci. 2002 Dec 1;22(23):10067–71.

17. Vazdarjanova A, Ramirez-Amaya V, Insel N, Plummer TK, Rosi S, Chowdhury S, et al. Spatial exploration induces ARC, a plasticity-related immediate-early gene, only in calcium/calmodulin-dependent protein kinase II-positive principal excitatory and inhibitory neurons of the rat forebrain. J Comp Neurol. 2006 Sep 20;498(3):317–29.

18. Bottai D, Guzowski JF, Schwarz MK, Kang SH, Xiao B, Lanahan A, et al. Synaptic activity-induced conversion of intronic to exonic sequence in Homer 1 immediate early gene expression. J Neurosci Off J Soc Neurosci. 2002 Jan 1;22(1):167–75.

19. Bramham CR, Worley PF, Moore MJ, Guzowski JF. The immediate early gene arc/arg3.1: regulation, mechanisms, and function. J Neurosci Off J Soc Neurosci. 2008 Nov 12;28(46):11760–7.

20. Bramham CR, Alme MN, Bittins M, Kuipers SD, Nair RR, Pai B, et al. The Arc of synaptic memory. Exp Brain Res Exp Hirnforsch Expérimentation Cérébrale. 2010 Jan;200(2):125–40.

21. Steward O, Worley PF. A cellular mechanism for targeting newly synthesized Arc mRNA to active synapses requires NMDA receptor activation. Neuron. 2001;30:227–40.

22. Guzowski JF, Miyashita T, Chawla MK, Sanderson J, Maes LI, Houston FP, et al. Recent behavioral history modifies coupling between cell activity and Arc gene transcription in hippocampal CA1 neurons. Proc Natl Acad Sci U S A. 2006 Jan 24;103(4):1077–82.

23. Carpenter-Hyland EP, Plummer TK, Vazdarjanova A, Blake DT. Arc expression and neuroplasticity in primary auditory cortex during initial learning are inversely related to neural activity. Proc Natl Acad Sci U S A. 2010 Aug 17;107(33):14828–32.

24. Ramirez-Amaya V, Vazdarjanova A, Mikhael D, Rosi S, Worley PF, Barnes CA. Spatial exploration-induced Arc mRNA and protein expression: evidence for selective, network-specific reactivation. J Neurosci. 2005;25:1761–8.

25. Szumlinski KK, Kalivas PW, Worley PF. Homer proteins: implications for neuropsychiatric disorders. Curr Opin Neurobiol. 2006;16:251–7.

26. Fagni L, Worley PF, Ango F. Homer as both a scaffold and transduction molecule. Sci STKE. 2002;2002:re8-re8-re8–re8.

27. Nalloor R, Bunting KM, Vazdarjanova A. Encoding of Emotion-Paired Spatial Stimuli in the Rodent Hippocampus. Front Behav Neurosci. 2012 Jun 14;6:27.

28. Chawla MK, Lin G, Olson K, Vazdarjanova A, Burke SN, McNaughton BL, et al. 3D-catFISH: a system for automated quantitative three-dimensional compartmental analysis of temporal gene transcription activity imaged by fluorescence in situ hybridization. J Neurosci Methods. 2004;139:13–24.

29. Heisler JM, Morales J, Donegan JJ, Jett JD, Redus L, O’Connor JC. The attentional set shifting task: a measure of cognitive flexibility in mice. J Vis Exp JoVE. 2015 Feb 4;(96):51944.

30. Richard A. Britten, Arriyam Fesshaye, Alyssa Tidmore, Aiyi Liu, Ashley A. Blackwell. Loss of Cognitive Flexibility Practice Effects in Female Rats Exposed to Simulated Space Radiation. Radiat Res. 2023 Aug 1;200(3):256–65.

31. Bissonette GB, Powell EM, Roesch MR. Neural structures underlying set-shifting: roles of medial prefrontal cortex and anterior cingulate cortex. Behav Brain Res. 2013 Aug 1;250:91– 101.

32. Birrell JM, Brown VJ. Medial Frontal Cortex Mediates Perceptual Attentional Set Shifting in the Rat. J Neurosci. 2000 Jun 1;20(11):4320.

33. Chen HY, Yang CY, Hsieh TH, Peng CW, Chuang LL, Chang YL, et al. Effects of transcranial direct current stimulation on improving performance of delayed-reinforcement attentional set-shifting tasks in attention-deficit/hyperactivity disorder rat model. Behav Brain Res. 2023 Feb 2;437:114145.

34. Guzowski JF, Worley PF. Cellular compartment analysis of temporal activity by fluorescent in situ hybridization (catFISH). Curr Protoc Neurosci. 2001;1.8.1-1.8.16-1.8.1-1.8.16-1.8.1-1.8.16-1.8.1-1.8.16.

35. Dixon-Melvin R, Shanazz K, Nalloor R, Bunting KM, Vazdarjanova A. Emotional state alters encoding of long-term spatial episodic memory. Neurobiol Learn Mem. 2022 Jan;187:107562.

36. Paxinos G, Watson C. The Rat Brain in Stereotaxic Coordinates: Hard Cover Edition. Elsevier; 2006. 451 p.

37. West MJ. Stereological methods for estimating the total number of neurons and synapses: issues of precision and bias. Trends Neurosci. 1999 Feb 1;22(2):51–61.

38. Giustino TF, Maren S. The Role of the Medial Prefrontal Cortex in the Conditioning and Extinction of Fear. Front Behav Neurosci. 2015;9:298.

39. Capuzzo G, Floresco SB. Prelimbic and Infralimbic Prefrontal Regulation of Active and Inhibitory Avoidance and Reward-Seeking. J Neurosci Off J Soc Neurosci. 2020/05/11 ed. 2020 Jun 10;40(24):4773–87.

40. van Aerde KI, Heistek TS, Mansvelder HD. Prelimbic and infralimbic prefrontal cortex interact during fast network oscillations. PloS One. 2008 Jul 16;3(7):e2725–e2725.

41. Marek R, Xu L, Sullivan RKP, Sah P. Excitatory connections between the prelimbic and infralimbic medial prefrontal cortex show a role for the prelimbic cortex in fear extinction. Nat Neurosci. 2018 May 1;21(5):654–8.

42. Hoover WB, Vertes RP. Anatomical analysis of afferent projections to the medial prefrontal cortex in the rat. Brain Struct Funct. 2007;212:149–79.

43. Sánchez-Bellot C, AlSubaie R, Mishchanchuk K, Wee RWS, MacAskill AF. Two opposing hippocampus to prefrontal cortex pathways for the control of approach and avoidance behaviour. Nat Commun. 2022 Jan 17;13(1):339–339.

44. Maren S, Phan KL, Liberzon I. The contextual brain: implications for fear conditioning, extinction and psychopathology. Nat Rev Neurosci. 2013/05/02 ed. 2013 Jun;14(6):417–28.

45. Quirk GJ, Mueller D. Neural Mechanisms of Extinction Learning and Retrieval. Neuropsychopharmacology. 2007;33:56–72.

46. Pitman RK, Shin LM, Rauch SL. Investigating the pathogenesis of posttraumatic stress disorder with neuroimaging. J Clin Psychiatry. 2001;62(Suppl17):47–54.

47. Bremner JD. Functional neuroimaging in post-traumatic stress disorder. Expert Rev Neurother. 2007 Apr 1;7(4):393–405.

48. Barense MD, Fox MT, Baxter MG. Aged rats are impaired on an attentional set-shifting task sensitive to medial frontal cortex damage in young rats. Learn Mem. 2002;9(4):191–201.

49. Cernotova D, Stuchlik A, Svoboda J. Roles of the ventral hippocampus and medial prefrontal cortex in spatial reversal learning and attentional set-shifting. Neurobiol Learn Mem. 2021 Sep 1;183:107477.

50. Monk CS, Zhuang J, Curtis WJ, Ofenloch IT, Tottenham N, Nelson CA, et al. Human hippocampal activation in the delayed matching-and nonmatching-to-sample memory tasks: An event-related functional MRI approach. Behav Neurosci. 2002;116(4):716–21.

51. Daniel TA, Katz JS, Robinson JL. Delayed match-to-sample in working memory: A BrainMap meta-analysis. Biol Psychol. 2016 Oct 1;120:10–20.

52. Habeck C, Rakitin BC, Moeller J, Scarmeas N, Zarahn E, Brown T, et al. An event-related fMRI study of the neural networks underlying the encoding, maintenance, and retrieval phase in a delayed-match-to-sample task. Cogn Brain Res. 2005 May 1;23(2):207–20.

53. Schon K, Tinaz S, Somers DC, Stern CE. Delayed match to object or place: An event-related fMRI study of short-term stimulus maintenance and the role of stimulus pre-exposure. NeuroImage. 2008 Jan 15;39(2):857–72.

